## Supplementary Information for "HYDRA: Fabrication of cell culture HYDrogels by Robotic liquid handling Automation for high-throughput drug testing"

1. E. Torchia, M. Di Sante, B. Horda, J. Zimmermann, M. Pezzotti, A. Enrico, F. S. Pasqualini
   Synthetic Physiology Lab, Department of Civil Engineering and Architecture, University of Pavia, 27100, Pavia, Italy
2. M. Mihajlovic, J. U. Lind
   Department of Health Technology, Technical University of Denmark, 2800 Kgs. Lyngby, Denmark
3. E. Cimetta
   Department of Industrial Engineering, University of Padua, 35131 Padova, Italy
4. S. Gabriele
   Mechanobiology & Biomaterials Group, Research Institute for Biosciences, University of Mons, CIRMAP, 20 Place du Parc, Mons B-7000, Belgium
5. F. Auricchio
   Group of Computational Mechanics and Advanced Materials, Department of Civil Engineering and Architecture, University of Pavia, 27100, Pavia, Italy

This file includes:

- Fig. S1 to S13
- Supplementary Movie captions S1 to S9
- Supplementary Table S1

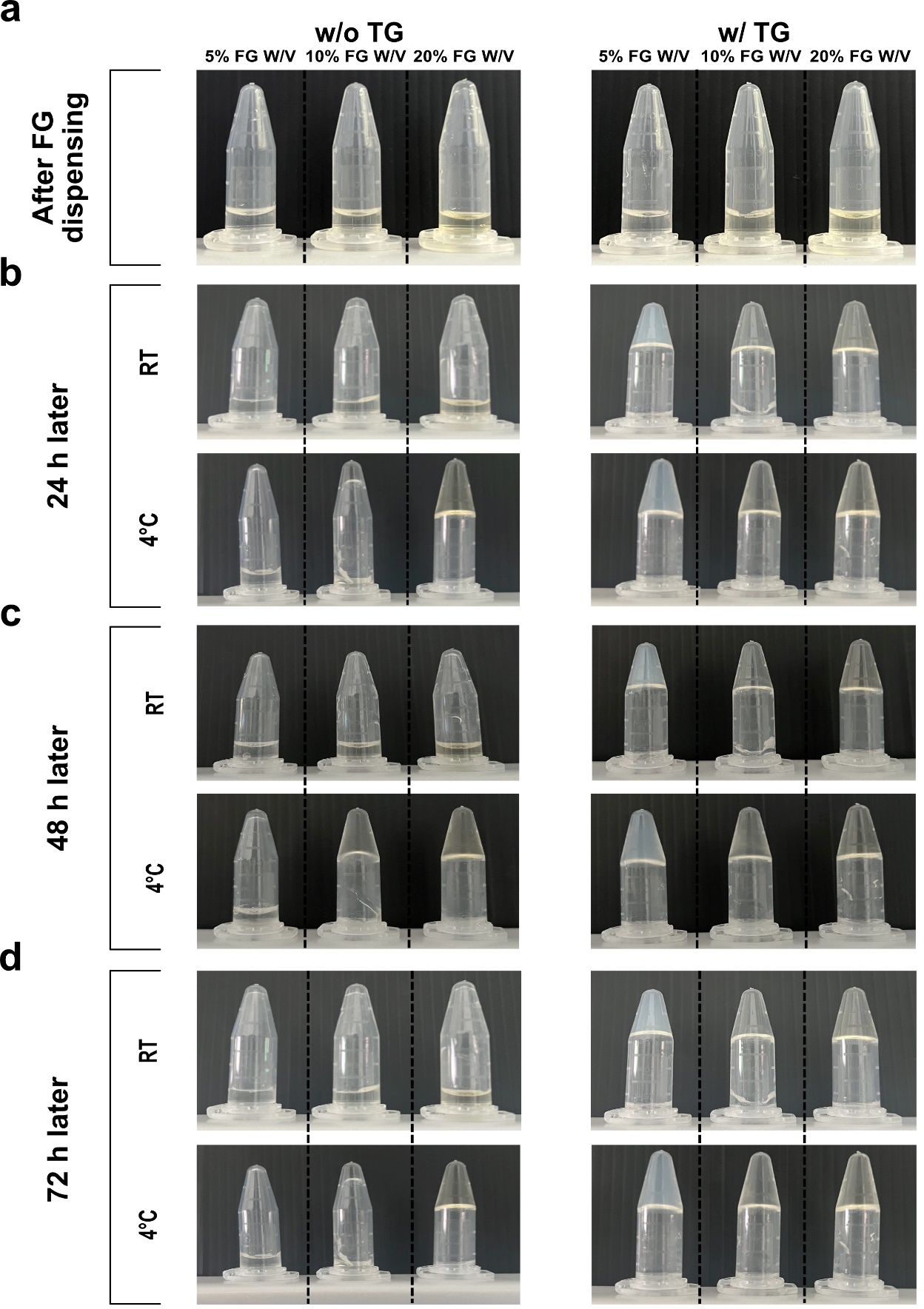

**Fig. S1. Detailed tube inversion analysis.** Photographic images (cross-sectional view) of FG mixtures in Eppendorf tubes with and without TG at room temperature (RT) and 4°C. Snapshots were taken (**a**) right after mixing, and every 24 h for 72 h **(b-c-d**).

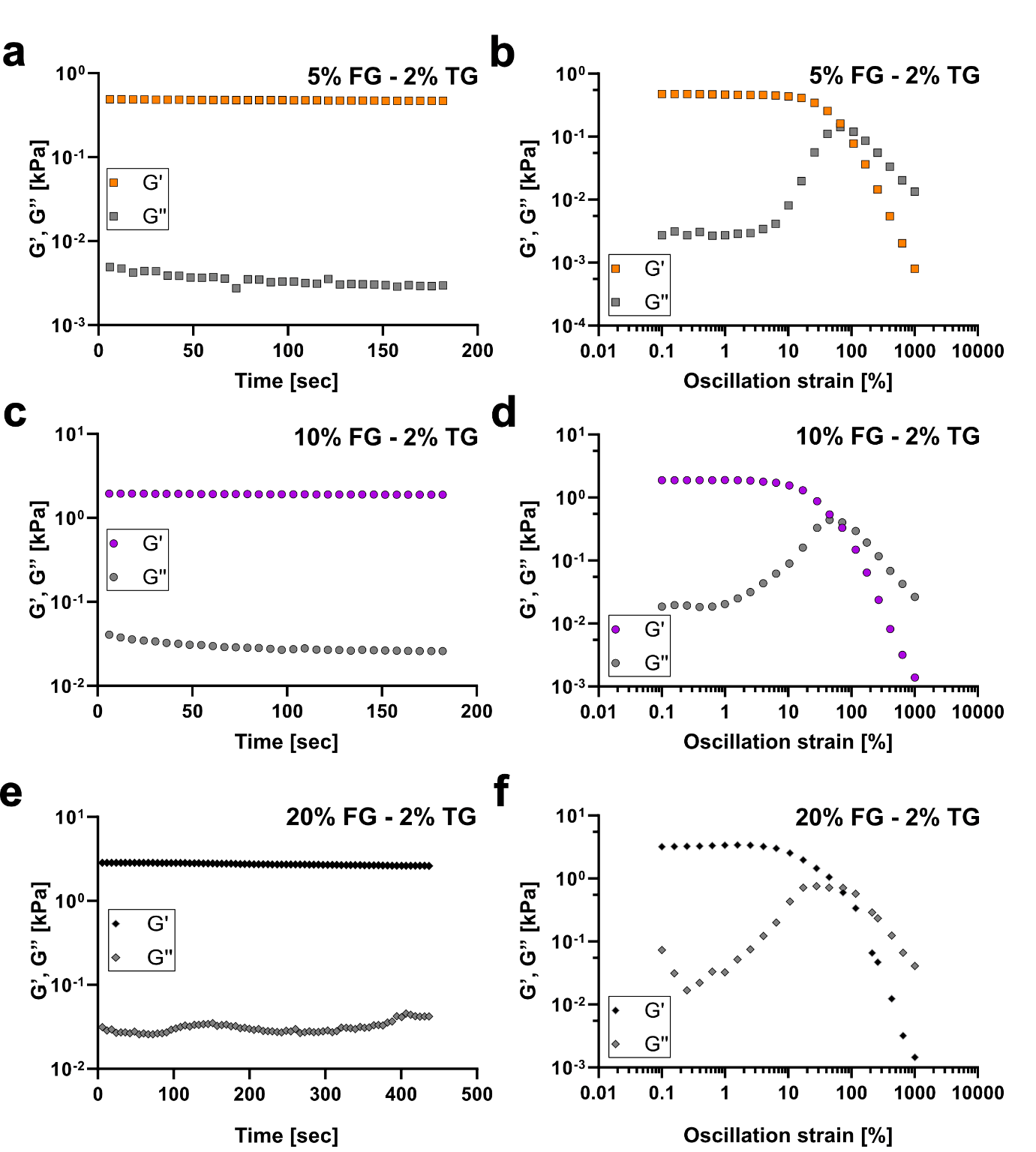
**Fig. S2. Fish gelatin hydrogels rheology.** Three FG concentrations (**a-b** 5%, **c-d** 10%, **e-f** 20% w/v) were considered while the TG concentration was fixed at 2% w/v. Storage (G’) and Loss Modulus (G’’) vs. time (sec) and oscillation strain (%). Measurements were performed on n=3 independently prepared formulations and kept at 37 °C. Samples were prepared one day before the measurements, after overnight crosslinking at 37 °C.

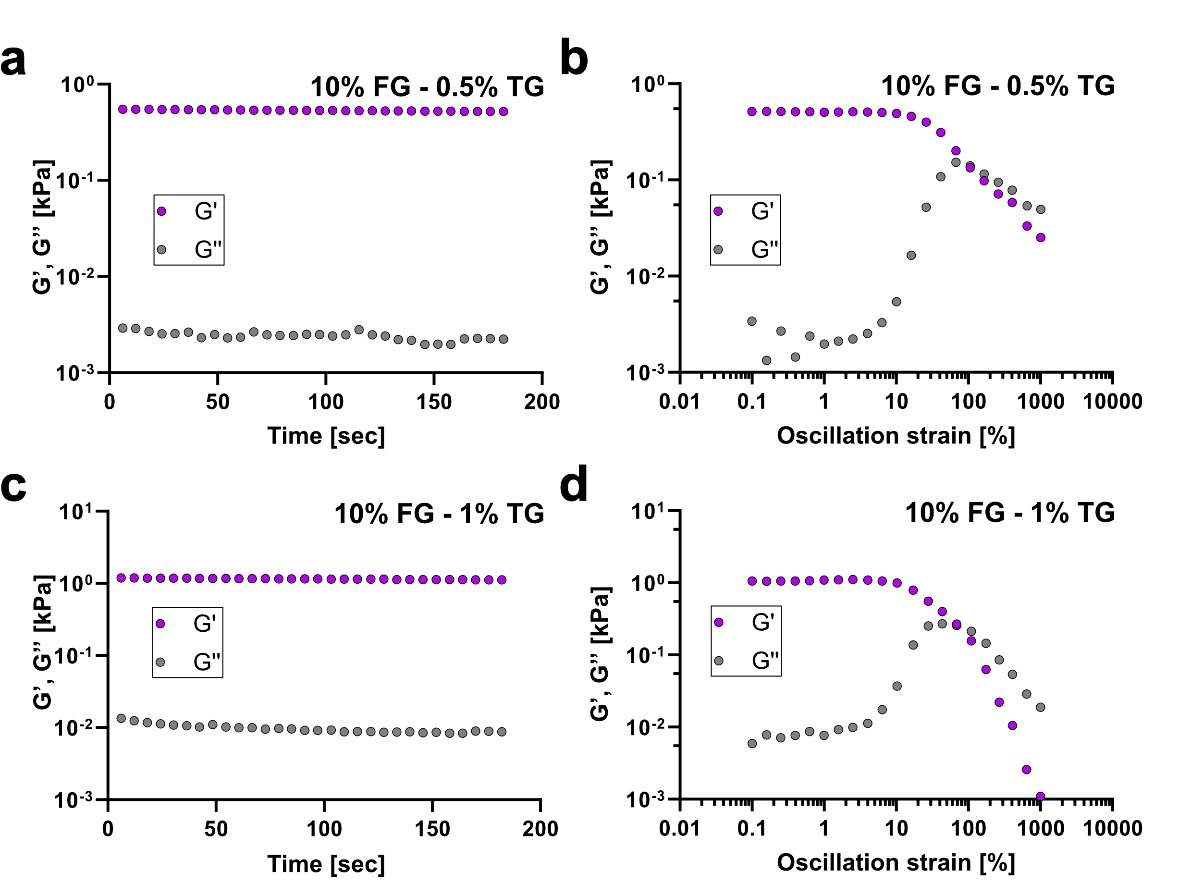

**Fig. S3. Fish gelatin hydrogels rheology.** Two TG concentrations (**a-b** 0.5%, **c-d** 1%) were considered while the FG concentration was fixed at 10% w/v. Storage (G’) and Loss Modulus (G’’) vs. time (sec) and oscillation strain (%). Measurements were performed on n=3 independently prepared formulations and kept at 37 °C. Samples were prepared one day before the measurements, after overnight crosslinking at 37 °C.

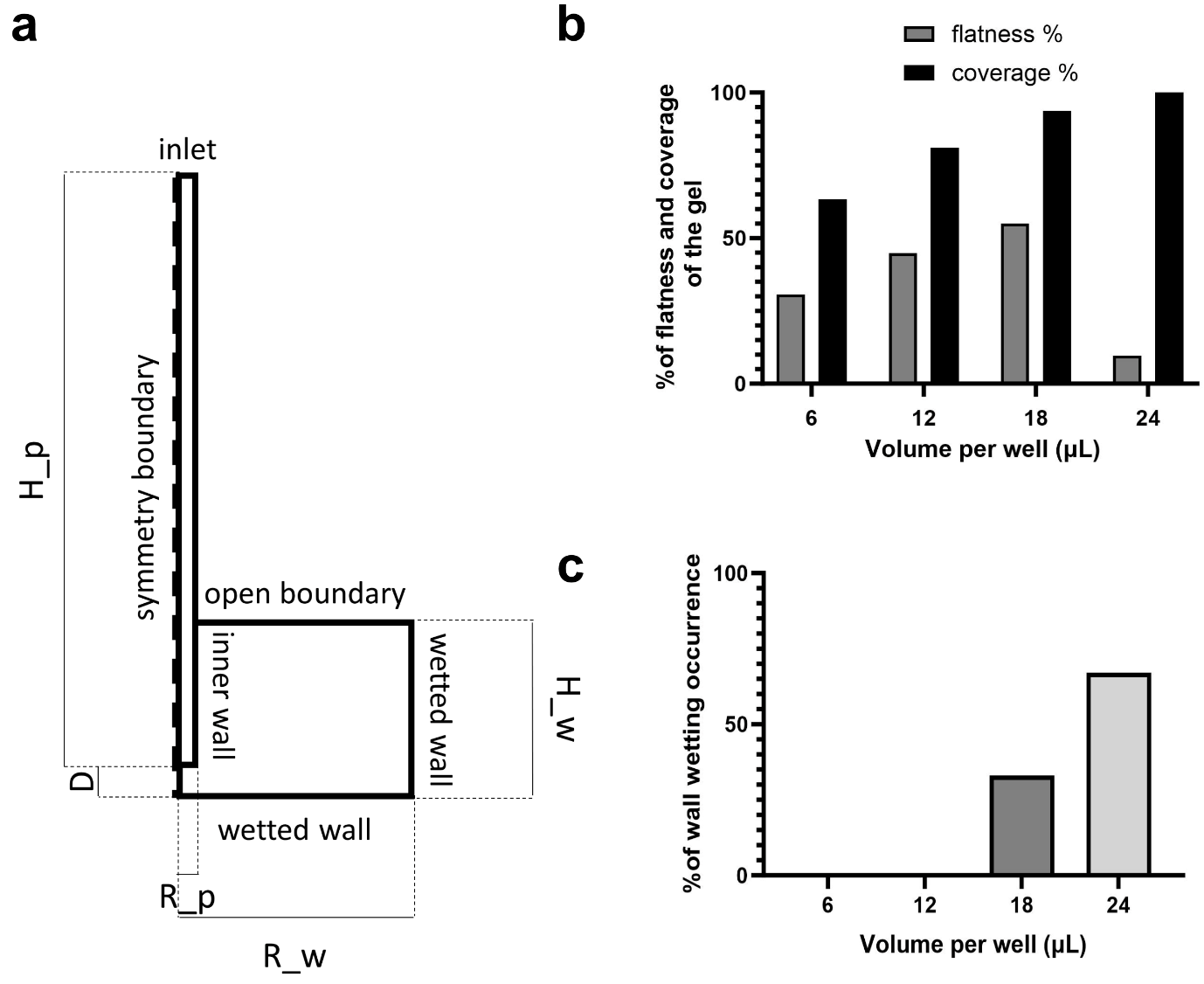

**Fig. S4. Modelling of the HYDRA dispensing method.** (**a**) Schematic representation of the geometry and boundary conditions of the COMSOL model. A 2D axisymmetric model was chosen to save computational power and time. The well is modelled as a rectangle of dimensions R_w and H_w, and the pipette is modelled as a rectangle of dimensions R_p and H_p. An inlet boundary condition is applied at the upper edge of the pipette, and an open boundary condition is applied at the upper edge of the well since it is open. Two wet wall boundary conditions are applied at the two walls of the well. (**b**) Flatness and coverage percentages of the gel concerning the radius of the well, evaluated for different dispensed volumes (6 µl, 12 µl, 18 µl, and 24 µl), using computer modelling. (**c**) The occurrence of wall wetting for different volumes using COMSOL simulations. For each nominal volume, nine combinations of values were considered, namely accounting for small variations of volume (±20%) and well radius (-100 µm, and -200 µm, which are respectively the single contributions of plate tolerance dimension and robot calibration, and the sum of them). The value of wall-wetting occurrence is calculated as the ratio of combinations of values in the sweep leading to wall-wetting.

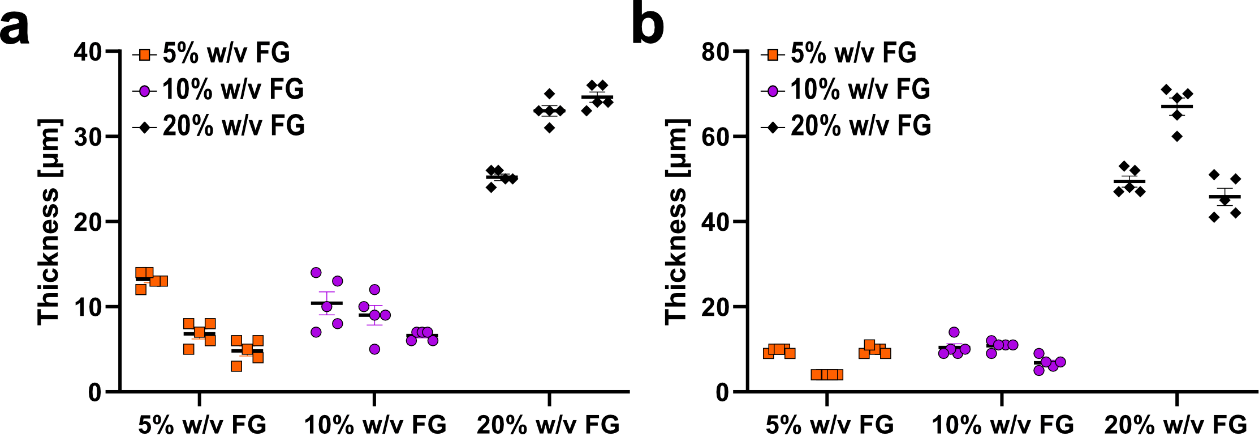

**Fig. S5. Assessment of thickness and flatness of fish gelatin (FG) hydrogels.** The graphs show two independent experiments (**a**-**b**) on hydrogel heights in the hydrated state (PBS 1x). Gel heights were calculated as the difference between the gel top (referenced using embedded fluorescent beads) and the gel bottom (referenced using HaCaT cell Actin on plastic). Images were obtained using confocal imaging. Three different concentrations were tested (5% - orange squares, 10% - purple dots, and 20% w/v – black diamonds). The data corresponds to single experiments (3 concentrations, n=5 samples for each concentration). For each sample, 5 FOVs were taken to measure the z-position and calculate thickness. Data are presented as mean ± s.e.m.

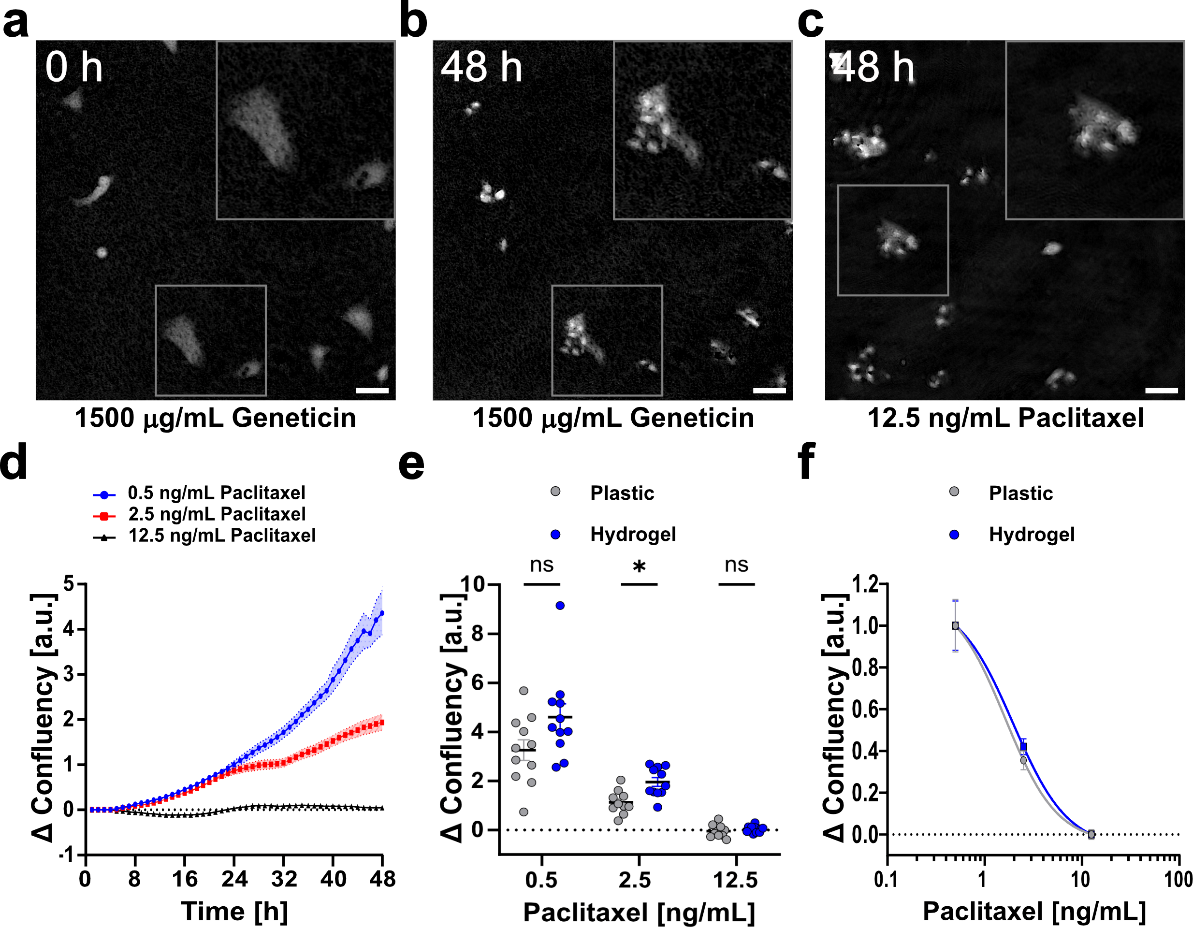

**Fig. S6. Drug test analysis using phase holographic imaging on hydrogel substrates.** (**a**) Holographic image of HaCaT after seeding (1500 μg mL^-1^ - positive control) and (**b**) 48 h after seeding. (**c**) Holographic image of HaCaT treated with 12.5 ng mL^-1^ of paclitaxel drug 48 h after seeding. Scale bars: 50 µm. (**d**) HaCaT cell confluency vs. time. Cells were cultured in paclitaxel-based cell culture media (0.5 ng mL^-1^ – blue dots, 2.5 ng mL^-1^ – red squares, 12.5 ng mL^-1^ black triangles) for 48 h on hydrogel substrates. Data were normalized concerning the initial confluency value. Solid lines represent mean values, and shaded areas represent s.e.m. (n=12). (**e**) Final cell confluency vs. paclitaxel concentration (0.5 ng mL^-1^, 2.5 ng mL^-1^, 12.5 ng mL^-1^) on plastic and hydrogel substrates. Data were normalized concerning the initial confluency value. (*) stands for significative difference. (**f**) Paclitaxel dose-response curves on plastic (IC50, 1.6 ng mL^-1^) and hydrogel (IC50, 1.8 ng mL^-1^) substrates (n=12). Data were normalized concerning the initial confluency value (n=12). Image brightness and contrast were adjusted for printed visibility.

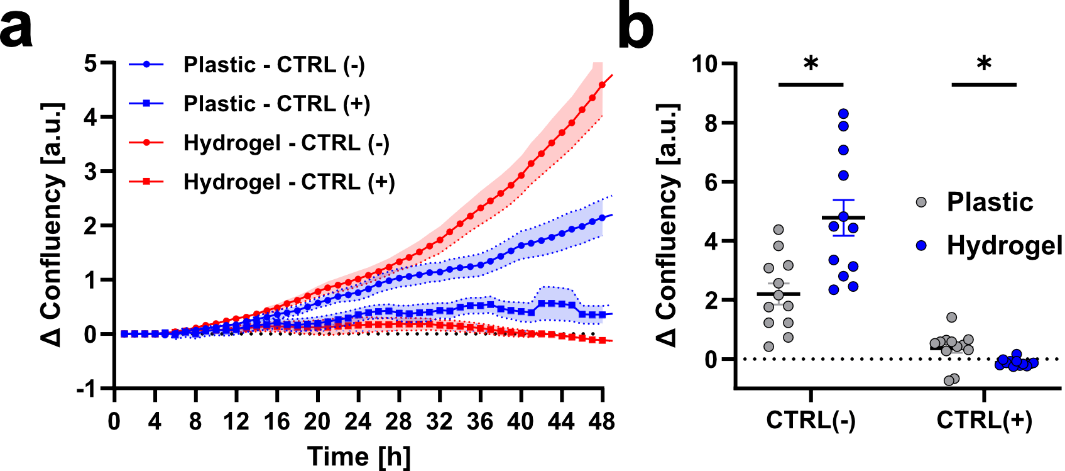

**Fig. S7. Control (CTRL) data from holographic imaging of nocodazole- and paclitaxel-treated cells on hydrogel and plastic.** (**a-b**) HaCaT cell confluency vs. time. Cells were cultured in 0.1% DMSO (CTRL (-)) or 1500 µg mL^-1^ Geneticin (CTRL (+)) on both hydrogel and plastic substrates for 48 h. Data were normalized concerning the initial confluency value. Solid lines represent mean values, and shaded areas represent s.e.m. (n=12). **B)** Final cell confluency on plastic and hydrogel substrates. Data were normalized concerning the initial confluency value (n=12). (*) stands for significative difference.

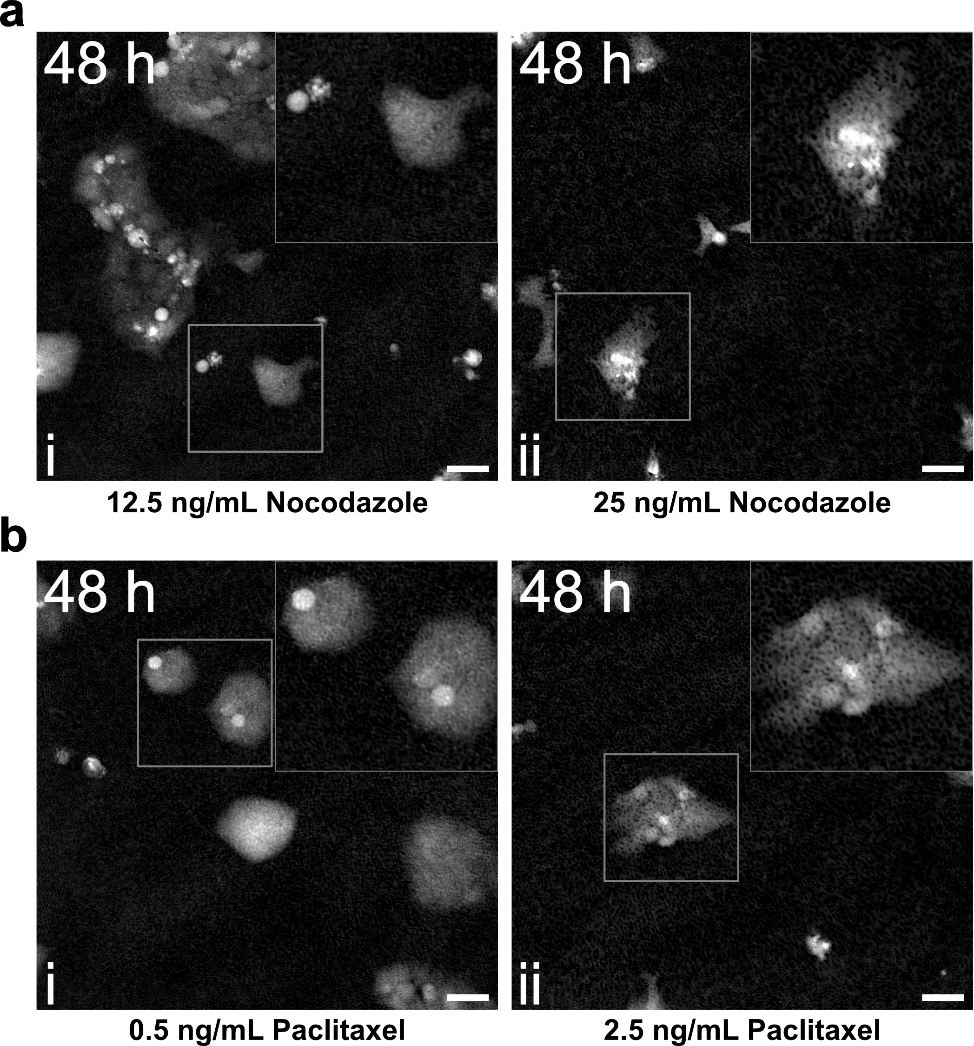

**Fig. S8. Holographic imaging of nocodazole- and paclitaxel-treated cells on hydrogel substrates.** (**a**) Holographic image of HaCaT treated with **(i)** 12.5 ng mL^-1^ and **(ii)** 25 ng mL^-1^ of nocodazole drug 48 h after seeding. (**b**) Holographic image of HaCaT treated with **(i)** 0.5 ng mL^-1^ and **(ii)** 2.5 ng mL^-1^ of paclitaxel drug 48 h after seeding. Image brightness and contrast were adjusted for printed visibility.

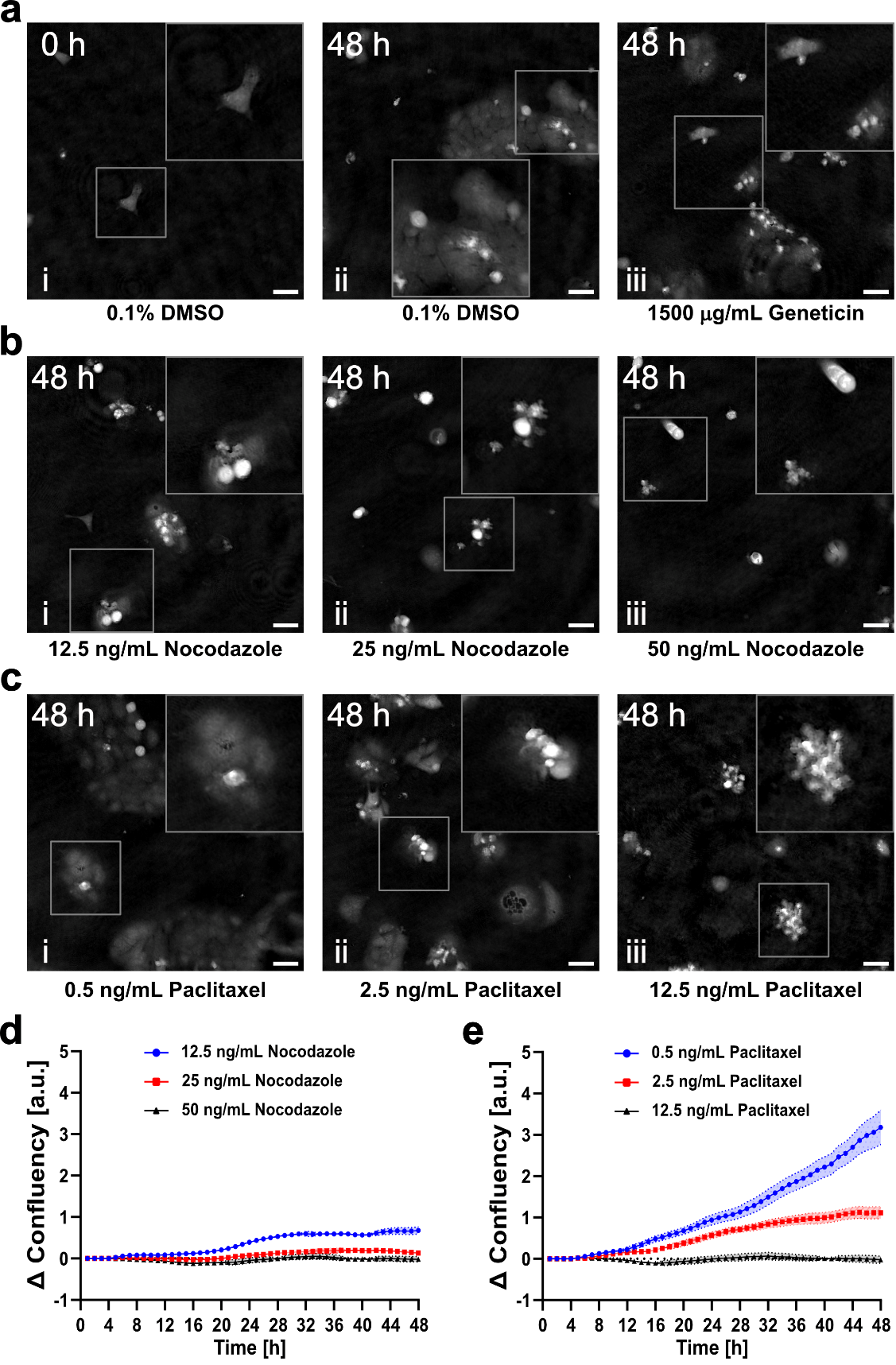

**Fig. S9. Holographic imaging analysis of nocodazole- and paclitaxel-treated cells on plastic substrates.** (**a**) Holographic image of HaCaT **(i)** after seeding (0.1% DMSO – vehicle negative control) and **(ii)** 48 h after seeding. **(iii)** Holographic image of HaCaT (1500 μg mL^-1^ - positive control) 48 h after seeding. (**b**) Holographic image of HaCaT treated with **(i)** 12.5 ng mL^-1^, **(ii)** 25 ng mL^-1^, **(iii)** and 50 ng mL^-1^ of nocodazole drug, 48 h after seeding. (**c**) Holographic image of HaCaT treated with **(i)** 0.5 ng mL^-1^, **(ii)** 2.5 ng mL^-1^, **(iii)** and 12.5 ng mL^-1^ of paclitaxel drug 48 h after seeding. (**d**) HaCaT cell confluency vs. time. Cells were cultured in nocodazole-based (12.5 ng mL^-1^ – blue dots, 25 ng mL^-1^ – red squares, 50 ng mL^-1^ black triangles) or (**e**) paclitaxel-based cell culture media (0.5 ng mL^-1^ – blue dots, 2.5 ng mL^-1^ – red squares, 12.5 ng mL^-1^ black triangles) for 48 h on plastic substrates. Data were normalized concerning the initial confluency value. Solid lines represent mean values, and shaded areas represent s.e.m. (n=12). Image brightness and contrast were adjusted for printed visibility.

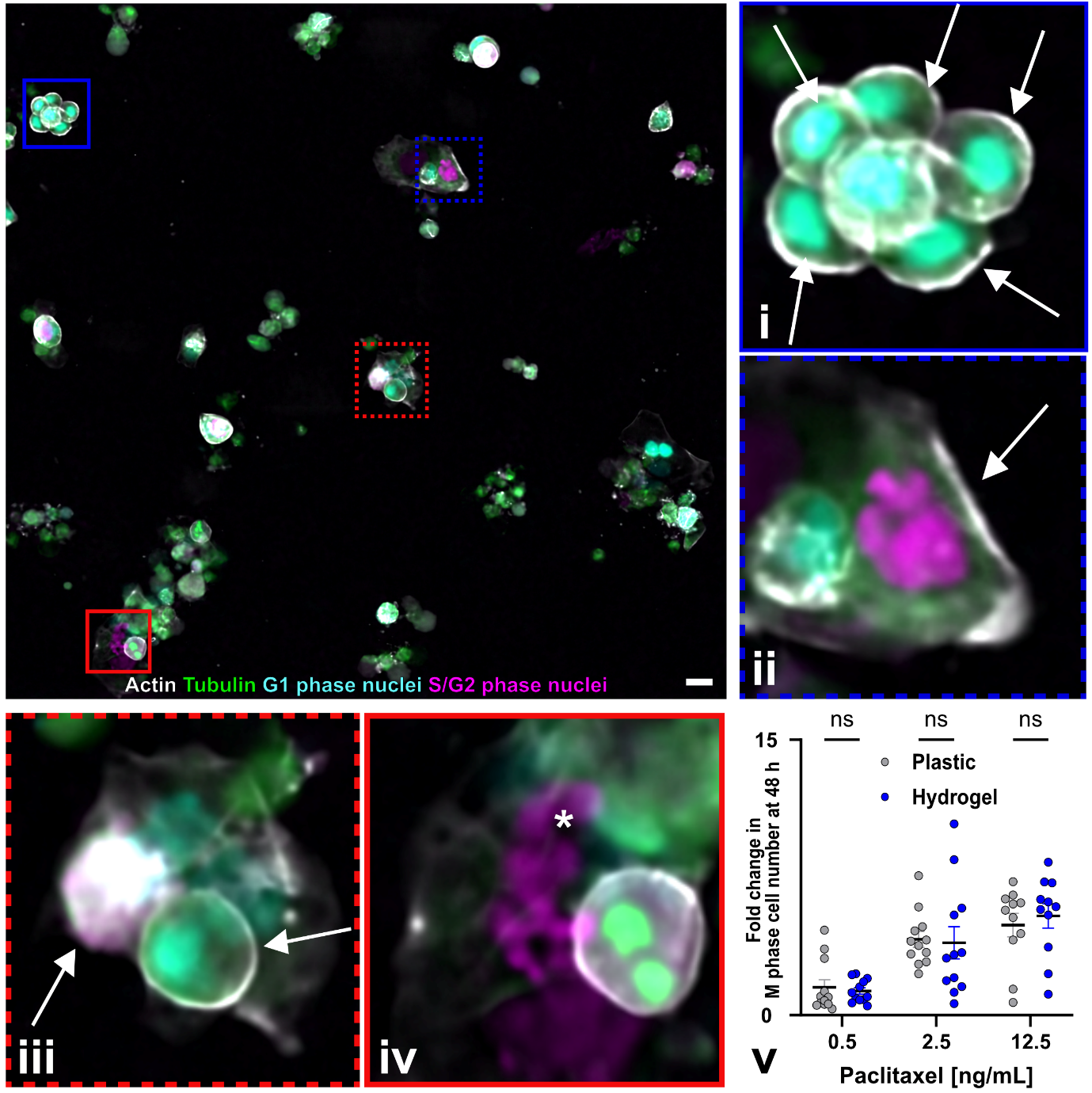

**Fig. S10. Drug test using fluorescence imaging on HYDRA hydrogels cast on inexpensive traditional tissue culture plastic.** Static widefield images of HaCaT (RFP – Life Act, in gray, GFP – tagged tubulin, in green) 48 h after seeding. Cells were treated with 12.5 ng mL^-1^ paclitaxel. Arrows stand for **(i)** Cells in the G1 phase (in cyan), **(ii)** cells in the S/G2 phase (in magenta), and **(iii)** cells in the mitotic (M) phase. The asterisk stands for **(iv)** nuclear fragmentation. **(v)** Fold change in M phase cell number after 48 h vs. paclitaxel concentration on plastic and hydrogel substrates. Cell number in the M phase was counted as a fraction of the total cell number in a FOV and then normalized to the vehicle negative control (n=12). Data are displayed as mean ± s.e.m. (ns) stands for no statistical difference. Images were taken on hydrogel thin layers cast on plastic plates. Scale bar: 25 μm.

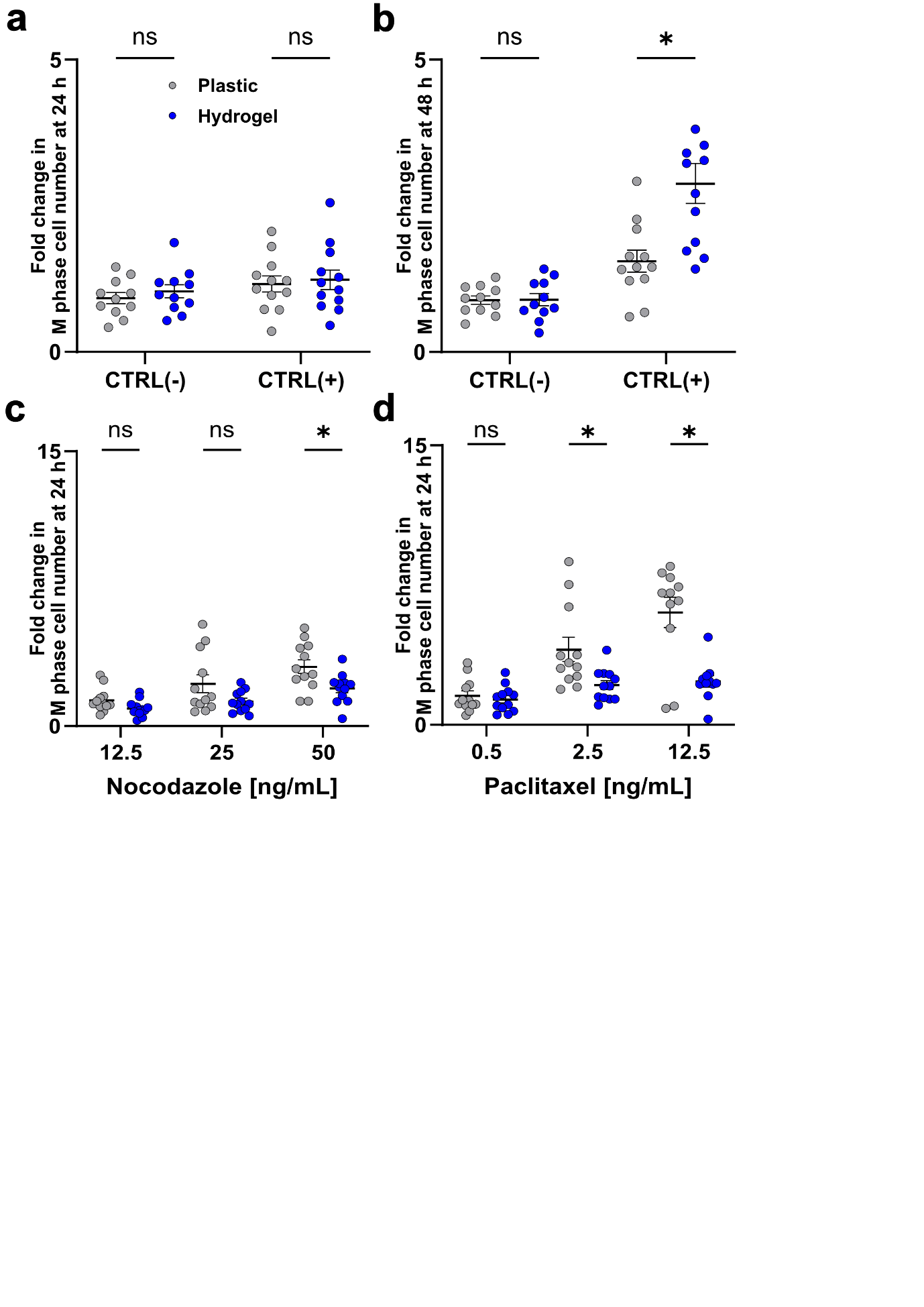

**Fig. S11. Static fluorescence imaging analysis of HYDRA hydrogels cast on inexpensive traditional tissue culture plastic.** Fold change in M phase cell number after 48 h vs. (**a-b**) negative and positive controls, (**c**) nocodazole, (**d**) paclitaxel concentration on plastic and hydrogel substrates. Cell number in the M phase was counted as a fraction of the total cell number in a FOV and then normalized to the vehicle negative control (n=12). Data are displayed as mean ± s.e.m. (ns) stands for no statistical difference.

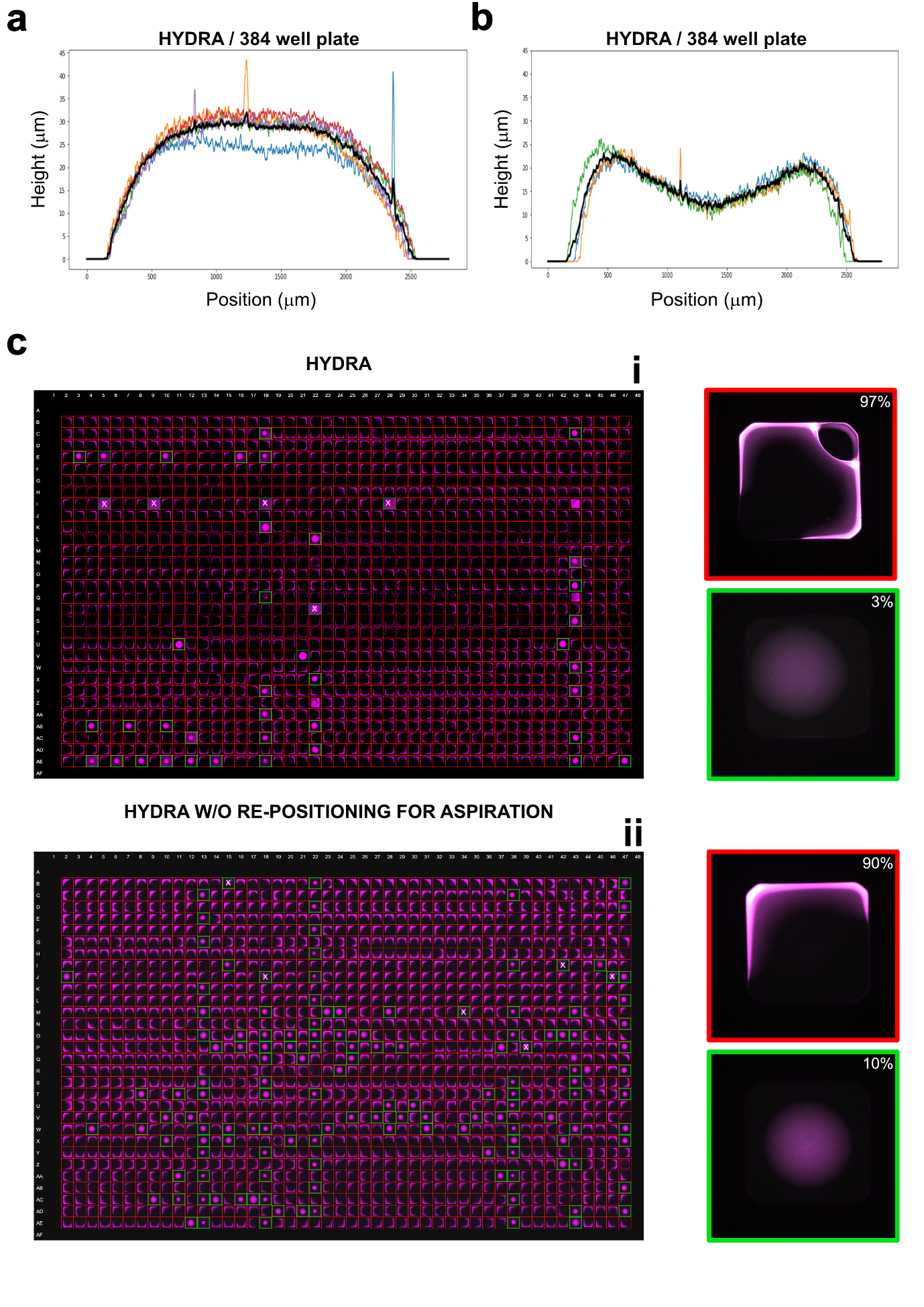

**Fig. S12. HYDRA-like hydrogels on 384- and 1536-well plates.** (**a**-**b**) Profile reconstructions of hydrogel samples in a 384-well plate showing two main populations: (**a**) one with a planar shape and (**b**) another with a concave shape. (**c**) HYDRA performed on a 1536-well plate. **i)** Using the HYDRA method, the hydrogels contact the well walls in almost the entire plate (97%) because the minimum liquid handling volume is too high for the well diameter. **ii)** Since robot movement between dispensing and aspiration could be a factor, if HYDRA is performed without repositioning for aspiration, the success rate increases slightly (from 3% to 10%). “x” symbols on well show wrong classification done by the automatic analysis.

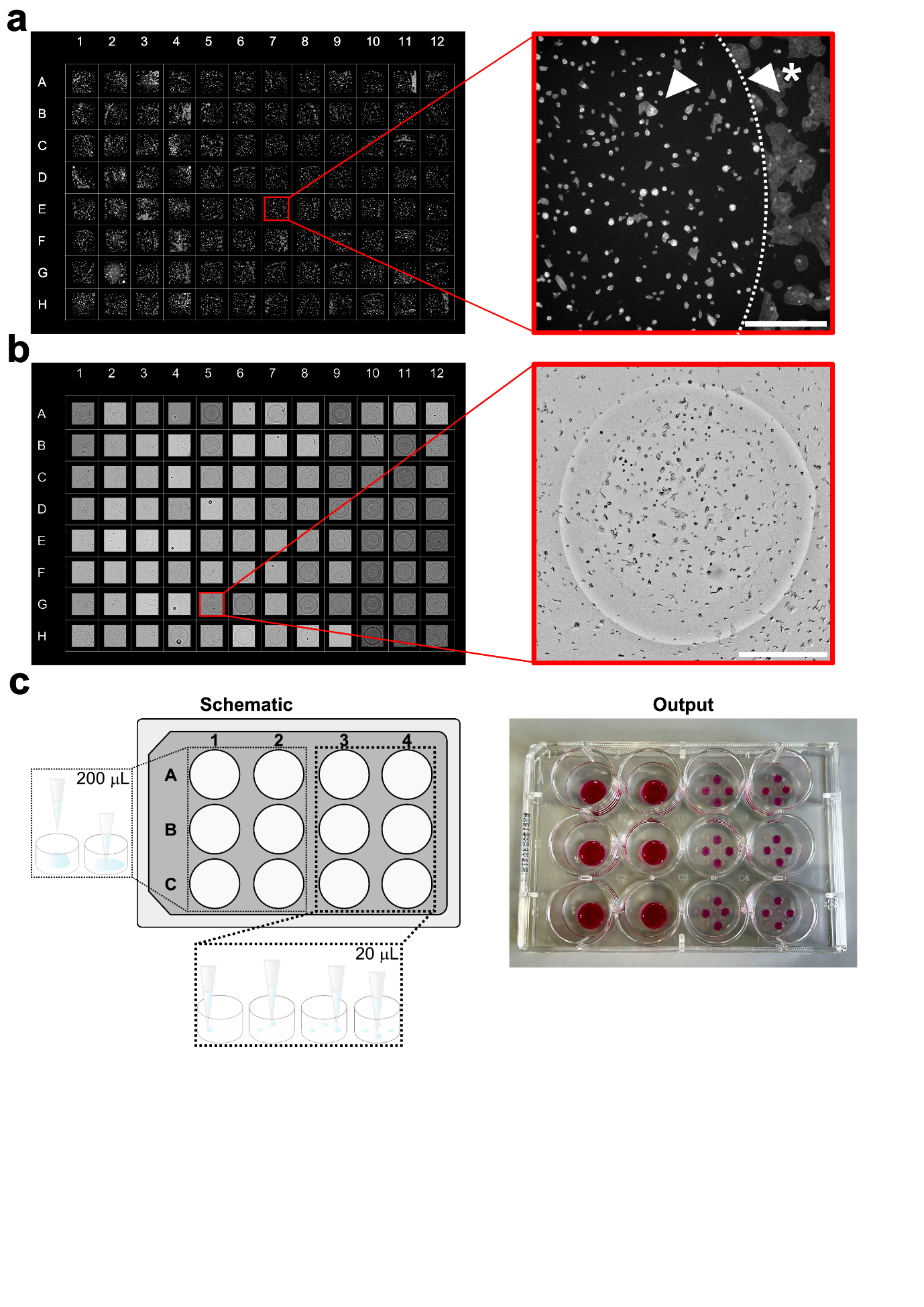

**Fig. S13. Further applications of HYDRA-like gels.** (**a**) 96-well plate tiling of HaCaT cells (actin in gray) seeded on hydrogels that were stored for two months in PBS at 4°C. Images were acquired with a 10x air objective (NA 0.3). Less than half of the hydrogel was captured (white arrow). A dashed line separates the hydrogel from the plastic substrate (white arrow with an asterisk). Scale bar: 500 μm. (**b**) 96-well plate tiling of HaCaT cells on hydrogels attached to glass. The plate with a glass bottom was first automatically functionalized using the OT-2 robot from Opentrons under the chemical hood, and then HYDRA was applied to cast the gels on the functionalized surface. To acquire the entire hydrogel with a 4x air objective (NA 0.13), 2 μL was used to cast the gels. Scale bar: 1 mm. (**c**) The HYDRA method can also be used in low-throughput plates. A schematic shows the custom protocol made using the OT-2 robot from Opentrons and a 12-well plate. The robot cast HYDRA-like hydrogels with a large volume (200 µL) in the first half of the plate, while increasing throughput in the second half by fabricating 4 HYDRA-like gels (20 µL each), which are 10 times smaller than the others. The output image shows the entire plate with hydrogel precursor solution pre-mixed with red food coloring (2 µL left to visualize the gels in this experiment).

**Supplementary Movie S1. Phase field simulation of large volume dispensing with meniscus formation.** Simulation of dispensing of a large volume of hydrogel precursor solution that touches the walls and forms the classic meniscus, using the *‘Laminar Two-phase Flow, Phase Field”* interface in COMSOL. In this Supplementary Movie, there is both the volume fraction of fluid (on the left) and the profile angle (on the right) visualization. In the volume fraction of fluid visualization, the two colors (blue and red) are used to represent the two different fluids, respectively hydrogel precursor solution and air, and the transition between the two colors represents the interface between the fluids. In the profile angle visualization, the colors represent the angle of the dispensed liquid at each point of the interface between the hydrogel precursor solution and air.

**Supplementary Movie S2. Phase field simulation of HYDRA dispensing method.** Simulation of dispensing and re-aspiration of 12 µl of hydrogel precursor solution, using the *‘Laminar Two-phase Flow, Phase Field”* interface in COMSOL. In this Supplementary Movie, there is both the volume fraction of fluid (on the left) and the profile angle (on the right) visualization. In the volume fraction of fluid visualization, the two colors (blue and red) are used to represent the two different fluids, respectively hydrogel precursor solution and air, and the transition between the two colors represents the interface between the fluids. In the profile angle visualization, the colors represent the angle of the dispensed liquid at each point of the interface between the hydrogel precursor solution and air.

**Supplementary Movie S3. Automated fabrication of hydrogel thin films in high throughput using a liquid handling system.** The Supplementary Movie shows the HYDRA method we use to cast hydrogel thin films in a 96-well plate by using a robot from Integra. The first protocol starts by mixing two solutions placed in the tube rack (on the right side of the field of view). Then, a second protocol uses the pre-mixed solution (in red) prepared before to dispense and re-aspirate the gels along columns, by placing the robot tip near the bottom of the 96-well plate (<=100 um). 20% w/v fish gelatin solution (with a red colorant) and 4% w/v transglutaminase were used in this Supplementary Movie. A volume of 1 µL of solution was intentionally retained to enhance the visualization of the hydrogel layers.

**Supplementary Movie S4. Automated fabrication of hydrogel thin films in high throughput using a liquid handling system: zoom on a single well of a 96-well plate.** The Supplementary Movie shows a close-up of a well in the 96-well plate using the HYDRA method. The pre-mixed hydrogel solution (in red) is dispensed and re-aspirated in the well, by placing the robot tip near the bottom of the 96-well plate (<=100 um). 20% w/v fish gelatin solution (with a red colorant) and 4% w/v transglutaminase pre-mixed solution were used in this Supplementary Movie. A volume of 1 µL of solution was intentionally retained to enhance the visualization of the hydrogel layer.

**Supplementary Movie S5. Automated holographic imaging of cells on hydrogel substrate.** The Supplementary Movie shows a series of 8 time-lapse sequences of the 96-well plate utilized for conducting a drug test. Each of the 8 fields of view (FOVs) corresponds to a specific condition of the drug test conducted on hydrogel substrates. These conditions are, as follows: the vehicle negative control (0.1% DMSO), positive control (1500 µg mL^-1^ geneticin), nocodazole concentrations of 12.5 ng mL^-1^, 25 ng mL^-1^, and 50 ng mL^-1^, as well as paclitaxel concentrations of 0.5 ng mL^-1^, 2.5 ng mL^-1^, and 12.5 ng mL^-1^. Scale bar: 50 μm.

**Supplementary Movie S6. Automated holographic imaging of cells on a plastic substrate.** The Supplementary Movie shows a series of 8 time-lapse sequences of the 96-well plate utilized for conducting a drug test. Each of the 8 fields of view (FOVs) corresponds to a specific condition of the drug test conducted on plastic substrates. These conditions are, as follows: the vehicle negative control (0.1% DMSO), positive control (1500 µg mL^-1^ geneticin), nocodazole concentrations of 12.5 ng mL^-1^, 25 ng mL^-1^, and 50 ng mL^-1^, as well as paclitaxel concentrations of 0.5 ng mL^-1^, 2.5 ng mL^-1^, and 12.5 ng mL^-1^. Scale bar: 50 μm.

**Supplementary Movie S7. HTS plate automatic acquisition.** The Supplementary Movie demonstrates the automated acquisition of HYDRA-like gels in a 384-well plate, with each field of view (FOV) representing a single well. This automated process is created using a Nikon JOB script and a 4X air objective (NA 0.13). For each well, the microscope moves to the center and captures a multi-channel image: one channel visualizes fluorescent beads embedded in the gel, and the other visualizes the well walls. This acquisition pipeline can be applied to every HTS plate by changing the corresponding plate in the script. The Supplementary Movie is accelerated by a factor of 10.

**Supplementary Movie S8. HTS plate analysis pipeline.** The Supplementary Movie shows the automatic quality control analysis of hydrogels in an HTS plate. The ImageJ macro script uses two channels acquired during the automatic acquisition: one visualizing beads embedded in the gel and the other visualizing the well walls. The script separates the two channels, applies thresholding, and generates a final composite image using a logical AND operation of the thresholded images. This composite image determines if the gels have contacted the walls. Additionally, the mean intensity of the mask obtained from the gel indicates whether the gel is planar or concave.

**Supplementary Movie S9. Automated long-term live fluorescence imaging on hydrogels.** The Supplementary Movie displays a long-term live fluorescence imaging of a cluster of HaCaT cells, genetically engineered to express fluorescent markers for Actin and our novel cell cycle sensor, FUCCIplex. In the Supplementary Movie, actin is visualized in gray, and nuclei, depending on their cell cycle phase, vary between cyan (G1 phase) and magenta (S/G2/M phase). The experiment was conducted using a 40X silicon oil objective (NA 1.25) with perfect focus on overtime. Images were taken every 15 min for 18 h. Scale bar: 25 μm.

**Supplementary Table S1. Parameters used in the COMSOL model.**

| **Parameter** | **Value** | **Unit** |
| --- | --- | --- |
| R_w | 3.45 | [mm] |
| H_w | 3 | [mm] |
| R_p | 0.16 | [mm] |
| H_p | 230 | [mm] |
| D | 0.3 | [mm] |
| Theta_adv | 40 | [°] |
| Theta_rec | 10 | [°] |
